# Frontoparietal pattern similarity analyses of cognitive control in monozygotic twins

**DOI:** 10.1101/2020.11.19.390492

**Authors:** Rongxiang Tang, Joset A. Etzel, Alexander Kizhner, Todd S. Braver

**Author notes:** **Corresponding Author:** Rongxiang Tang, One Brookings Drive, Campus Box 1125, Department of Psychological and Brain Sciences, Washington University in St. Louis, Saint Louis, Missouri, USA 63130.

## Abstract

The ability to flexibly adapt thoughts and actions in a goal-directed manner appears to rely on cognitive control mechanisms that are strongly impacted by individual differences. A powerful research strategy for investigating the nature of individual variation is to study monozygotic (identical) twins. Evidence of twin effects have been observed in prior behavioral and neuroimaging studies, yet within the domain of cognitive control, it remains to be demonstrated that the neural underpinnings of such effects are specific and reliable. Here, we utilize a multi-task, within-subjects event-related neuroimaging design with functional magnetic resonance imaging to investigate twin effects through multivariate pattern similarity analyses. We focus on fronto-parietal brain regions exhibiting consistently increased activation associated with cognitive control demands across four task domains: selective attention, context processing, multi-tasking, and working memory. Healthy young adult monozygotic twin pairs exhibited increased similarity of within- and cross-task activation patterns in these fronto-parietal regions, relative to unrelated pairs. Twin activation pattern similarity effects were clearest under high control demands, were not present in a set of task-unrelated parcels or due to anatomic similarity, and were primarily observed during the within-trial timepoints in which the control demands peaked. Together, these results indicate that twin similarity in the neural representation of cognitive control may be domain-general but also functionally and temporally specific in relation to the level of control demand. The findings suggest a genetic and/or environmental basis for individual variation in cognitive control function, and highlight the potential of twin-based neuroimaging designs for exploring heritability questions within this domain.

## 1. Introduction

Cognitive control refers to the ability to regulate, coordinate, and sequence thoughts and actions in accordance with internally maintained goals (Braver, 2012). Convergent theoretical and empirical work suggests that cognitive control is a domain-general capacity (Duncan, 2010), involving mental processes spanning multiple domains of cognition, with shared neural representations within the fronto-parietal network (FPN) (Braver et al., 2009; Miller & Cohen, 2001; Miyake et al., 2000). Impairment in cognitive control is often detrimental to human functioning (Braver et al., 2002; Synder et al., 2015; McTeague et al., 2016; Barch & Sheffield, 2017). Consequently, identifying sources of individual variation in cognitive control function is of both theoretical and translational importance, for example, as a first step towards identifying critical intervention targets.

Twin-based designs provide an opportunity to examine potential sources of individual variation in cognitive control: if cognitive control is heritable, then its variability should at least be partially explained by genetic and environmental relatedness. In particular, monozygotic twins should exhibit the highest levels of similarity, as they share 100% of segregated loci and similar environments (Polderman et al., 2015). Indeed, in a pioneering and influential behavioral study, using a twin-based multi-task design, individual differences in cognitive control function were shown to be almost entirely genetic in origin (Friedman et al., 2008). Based on these findings, it is important to determine whether the systematic nature of individual variation in cognitive control can also be demonstrated in terms of neural substrates. More recently, neuroimaging studies have begun to address this question, by examining the heritability of brain activity patterns within various domains of cognitive control. In a series of large-scale studies, Blokland and colleagues (2008, 2011, 2017) demonstrated significant heritability effects on brain activation patterns within the FPN during a working memory task (N-back). Several small-scale twin neuroimaging studies showed convergent evidence of heritability in working memory and executive control (Koten et al., 2009; Matthews et al., 2007).

Recently, a powerful approach that is starting to be used in twin neuroimaging studies is that of multivariate pattern similarity analysis. This approach enables computation of a clear and compact quantitative metric that indicates the degree of similarity in neural activation patterns between paired individuals. In particular, reliable twin activation similarity effects have been observed in functionally selective brain regions during visual tasks, such as the fusiform face area (Polk et al., 2007; Pinel et al., 2015). Utilizing the same approach, we recently demonstrated robust patterns of twin similarity in FPN activation patterns during N-back task performance in Human Connectome Project data (Etzel et al., 2020).

However, none of these prior neuroimaging studies demonstrated clear twin similarity within the domain of cognitive control, as they involved single tasks, and either focused on different domains, or did not selectively target cognitive control (e.g., N-back load manipulations may engage potentially different processes, such as working memory). Furthermore, as the prior studies utilized blocked experimental designs, they were unable to identify the temporal dynamics of twin similarity in neural activation patterns in relationship to task demands. Consequently, three key questions remain regarding twin similarity effects in neural activation patterns related to cognitive control. In particular, are these effects: 1) *domain-general (*or only observed in specific tasks, such as the N-back); 2) *anatomically specific* (i.e., preferentially observed in brain regions strongly associated with control demands); and 3) *temporally specific* (i.e., preferentially expressed during time periods of peak control demand)?

Here, we address these three questions by examining twin similarity in activation patterns across four prototypical task domains associated with cognitive control (selective attention, context processing, multi-tasking, and working memory), utilizing event-related designs and a task battery specifically developed for examining both intra- and inter-individual variation in cognitive control. We focused our examination of twin similarity on FPN brain parcels demonstrating consistent sensitivity to control demands in univariate contrasts in previous analyses (Braver et al., 2020). We then compared twin effects to those observed within a *null set* of parcels unengaged by cognitive control demands. Finally, we examined the time course of twin similarity effects within task trials to determine their potential temporal specificity. In all analyses, monozygotic twin pairs were compared to a set of matched unrelated pairs. The goal of these analyses was to provide an in-depth exploration of twin similarity as a window from which to understand the degree of specificity present in the neural basis of individual differences in cognitive control. For this reason, this first-stage of investigation did not include other sets of related pairs (e.g., dizygotic twins, siblings) since the focus was on the nature of individual differences rather than pure genetic modeling. Nevertheless, we return to this issue in the Discussion, suggesting how the approach pioneered here can be extended to address more direct questions regarding the degree to which the individual difference profiles observed here can be decomposed into genetic versus environmental contributions.

It is worth noting that our study is a complementary yet unique extension of the prior behavioral study by Friedman and colleagues (2008). We not only employed a distinct and theoretically-focused set of tasks specifically designed to tap into cognitive control, which enables a conceptual replication of the Friedman et al (2008) findings, but also included novel neural measures that are sensitive in capturing idiosyncratic neural representations of cognitive control and their temporal dynamics. As such, the addition of neuroimaging assessment in the current study provides critical information regarding the neural substrates of individual difference in cognitive control, which can further advance our understanding of how individual variability in cognitive control function might arise.

## 2. Materials and Methods

### 2.1 Participants

Healthy participants ranging from 18 to 45 years old were recruited from local communities for a multi-session behavioral and neuroimaging project – the Dual Mechanisms of Cognitive Control (DMCC) (Braver et al., 2020). As part of this project, twenty-eight pairs (56 participants) of monozygotic twins (46 females, mean age= 31.70 years, SD= 7.17) were recruited via the Human Connectome Project (Van Essen et al., 2013), Missouri Family Registry, and neighborhood flyers. Using all 56 twin participants and a group of 38 unrelated participants from the same DMCC project, we also generated 47 pairs of unrelated individuals (94 participants), who were of the same sex and within 2 years of age within each pair (62 females, mean age= 31.85 years, SD= 6.03). All participants were screened and excluded if they were diagnosed with mental illness, neurological trauma, or MRI safety contraindications. The full listing of exclusion criteria can be found at the following link (https://osf.io/6efv8/). Participants provided written informed consent and received $450 compensation (plus a $10 maximum task performance bonus) for completing all sessions. All study protocols were approved by the Institutional Review Board at Washington University.

### 2.2 Imaging Acquisition

All imaging data were acquired at Washington University in St. Louis on a Siemens 3T PRISMA scanner using a 32-channel head coil. High-resolution MPRAGE anatomical scans (T1- and T2-weighted, voxel size=0.8 mm isotropic, Repetition Time (TR) =2400 msec for T1 and 3200 msec for T2) and blood-oxygen-level-dependent (BOLD) functional scans (voxel size=2.4 mm isotropic, TR=1200 msec), were acquired using the multi-band accelerated echo-planar imaging pulse sequences (acceleration factor=4, alternating anterior to posterior and posterior to anterior phase-encoding directions, no in-plane acceleration) from the Center for Magnetic Resonance Research (CMRR) at University of Minnesota. Further details related to acquisition protocol and parameters can be found at the following link (https://osf.io/tbhfg/).

### 2.3 Experimental Design and Statistical Analyses

#### 2.3.1 Procedure and Task Design

All participants underwent four sessions (one behavioral, three neuroimaging) in the DMCC study. In the behavioral session, participants completed self-report measures of personality and psychological health, cognitive assessments of crystallized and fluid intelligence, and other relevant cognitive domains. In each of the three neuroimaging sessions, the same four tasks were performed, but with different experimental manipulations and conditions applied. The present analysis only uses data from the first (Baseline) neuroimaging session, as this session includes task variants specifically designed to capture individual differences in cognitive control, while also avoiding practice effects that might be present in the subsequent two sessions. Participants performed each task across two consecutive BOLD scanning runs (8 task runs total per session), each approximately 12 minutes in duration. The task order was randomized across participants, but to facilitate twin-based comparisons, the task order was the same for both members of a twin pair.

Participants were scanned while performing four well-established cognitive control tasks: the color-word Stroop task (Stroop), AX-Continuous Performance Test (AX-CPT), Cued Task-Switching (Cued-TS), and Sternberg Working Memory (Sternberg). In all tasks except the Stroop, participants responded with a manual response box inside the scanner, using index and middle fingers of the right hand. In the Stroop, participants made vocal responses that were recorded through a microphone. Prior to performing the tasks inside the scanner, participants practiced a short block of each task on a computer using a button box with the same key mapping as the scanner response box. For Stroop, they were asked to say each response out loud. All tasks were programmed and presented to participants using Eprime software (Version 2.0, Psychology Software Tools, Pittsburgh, PA). In the following section, we briefly describe each task and its key contrasts. Additional description of each task condition and associated theoretical rationale can be found in Braver et al. (2020); a detailed listing of all task stimuli and parameters can be found at the following link (https://osf.io/48aet/).

##### Stroop

The color-word Stroop is widely recognized as a canonical task of cognitive control (Stroop, 1935; Egner & Hirsch, 2005), in which top-down selective attention is required to focus processing on the task-relevant font color of printed words, while ignoring the irrelevant but otherwise dominant word name. In the current version of the task, participants were asked to name the font color of the word (red, purple, blue, yellow, white, pink, black or green) out loud. Because of the large number of different responses (8), the task was implemented with vocal rather than manual responding. The Stroop interference effect, which contrasts incongruent (word name indicates a different color than the font color, e.g., BLUE in red font) and congruent (word name matches font color, e.g., BLUE in blue font) trials (see Figure 1A) was the key contrast. We employed a proportion congruence (PC) manipulation (Bugg et al., 2008), such that high PC was used for a set of stimuli (i.e., biased trials). Under high PC conditions, congruent trials are frequent and incongruent trials are rare, producing robust Stroop interference and individual differences effects (Kane & Engle, 2003). In our task fMRI analysis, biased incongruent trials were assumed to have high cognitive control demands, while biased congruent trials had low demands.

**Figure 1.**
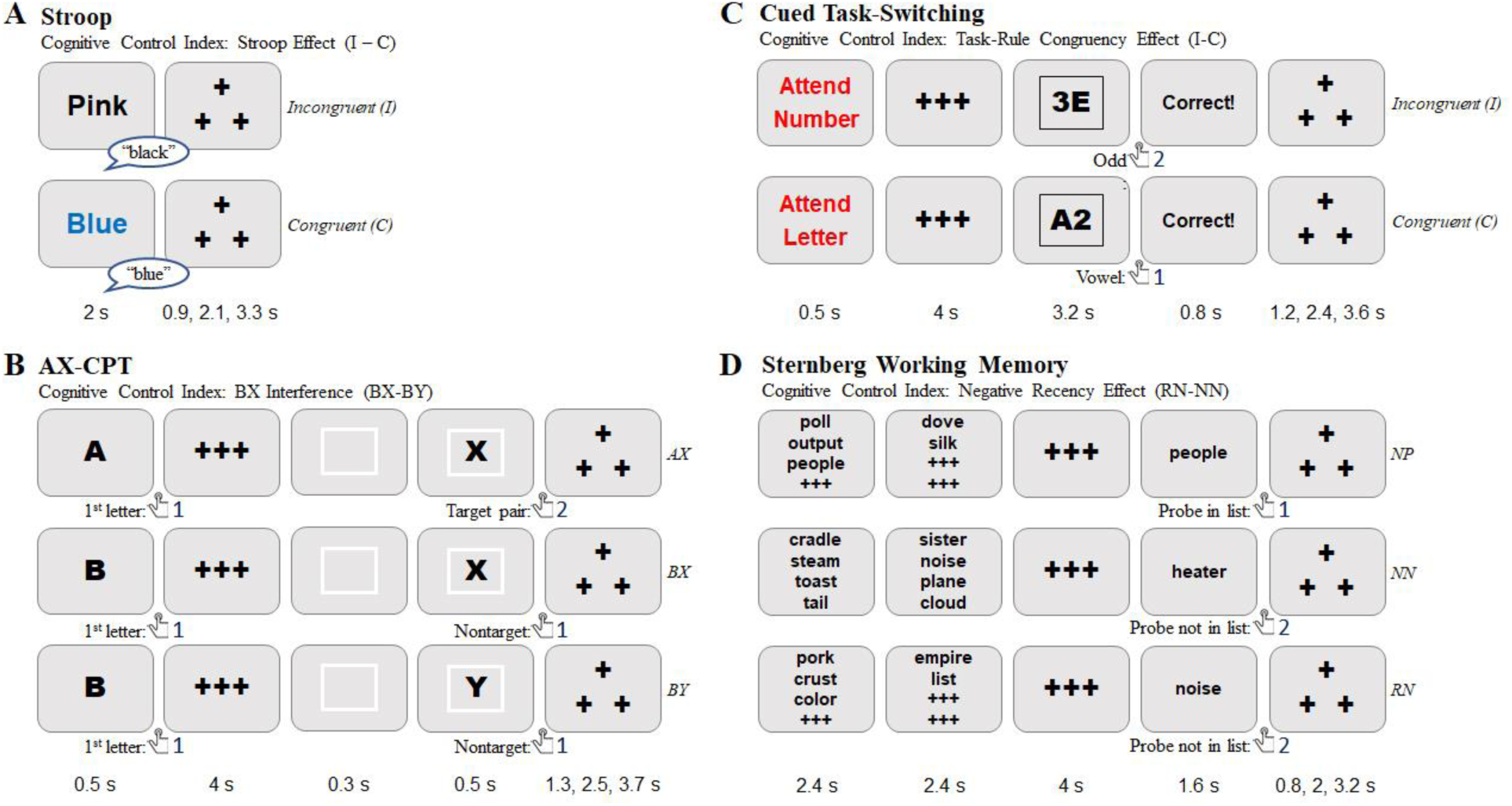
Diagram of Cognitive Control Tasks in DMCC battery

##### AX-CPT

The AX-CPT is increasingly used as a task of context processing and cognitive control, given its simplicity, flexibility and applicability in a wide-range of populations (Barch et al., 2008; Chatham et al., 2009; Chun et al., 2018; MacDonald, 2008; Paxton et al., 2007). In the task, participants were asked to respond to a pair of letters through button presses, with the first letter defined as a cue, followed by a second letter (sometimes a number), defined as a probe. The AX trials (A cue followed by an X probe) are the target response in the task, and occur with high frequency, leading to strong cue-probe associations. Cognitive control is postulated as a key process involved in maintaining and utilizing the contextual cue information, in order to minimize errors and response interference occurring on BX trials, which occur when the X-probe is presented but not preceded by an A-cue. Thus, a useful index of cognitive control for this task is the BX interference effect, which contrasts BX and BY (neither an A-cue nor an X-probe is presented, leading to low control demands) trial types (see Figure 1B). In our task fMRI analysis, BX trials were assumed to have high cognitive control demands, while BY trials had low demands.

##### Cued-TS

Cued task-switching (Cued-TS) has long been recognized as a critical paradigm to assess a core component of cognitive control – the ability to update and activate task-representations in an on-line manner to appropriately configure attention and action systems for processing the upcoming target (Kiesel et al., 2010; Meiran, 1996; Vandierendonck et al., 2010). In the task, participants were asked to respond to letter-digit target stimuli using button presses, with the target defined by the advance task cue (Letter or Digit). In the letter task, they were asked to decide if the letter is a consonant or a vowel. In the digit task, they were asked to decide if the digit is odd or even. An important index of cognitive control in task-switching paradigms is the task-rule congruency effect (TRCE; Meiran & Kessler, 2008), which refers to the increased interference (both errors and reaction time) when the response required for the current task trial is different than the response that would be required (to the same target stimulus) if the other task had been cued (see Figure 1C). In letter-digit task-switching (also called consonant-vowel, odd-even or CVOE; Minear & Shah, 2008; Rogers & Monsell, 1995), if a right button press is required for a consonant (left for a vowel) in the letter task, and a right button press for odd (left for even) in the digit task, then the “3E” target stimulus is incongruent and the “A2” target stimulus is congruent (since under either task, a left button press is correct). In this task, there were also two sets of stimuli. One set of stimuli were mostly congruent (80% congruent; 20% incongruent), whereas the other set were unbiased (50% congruent, 50% incongruent). Thus, to maximize individual differences, we used the biased set of stimuli in our task fMRI analysis, where incongruent trials were assumed to have high cognitive control demands, while congruent trials had low demands.

##### Sternberg

The Sternberg item-recognition task has been one of the most popular experimental paradigms used to assess short-term / working memory for over 50 years (Sternberg, 1966). In the task, participants were asked to remember a list of words (5-8 words) and then respond to a probe word, using a button press to indicate whether the probe word had been one of the words previously presented in the memory set for that trial. The Sternberg task has been adapted particularly for the study of cognitive control, and in neuroimaging paradigms, with the “recent probes” variant (Jonides et al., 1998; Jonides & Nee, 2006). In the recent probes variant, the key manipulation is that the probe item could have been a member of the memory set of the previous (not current) trial, which is termed a “recent negative” (RN) probe. On these RN trials, the probe is associated with high familiarity, which can increase response interference and errors, unless cognitive control is utilized to successfully determine that the familiarity is a misleading cue regarding probe status (target or nontarget). Thus, a key index of cognitive control in this Sternberg variant is the negative recency effect (Jonides & Nee, 2006; Monsell, 1978), which contrasts RN and NN trials (NN: novel negative, when the probe item is not a member of the current or previous trial’s memory set; see Figure 1D). The current task has two sets of memory set items. One is the critical set of 5 words, whereas the other is the high-load set of 6-8 words. To maintain the same load length, we focused our task fMRI analysis on the critical set (5-item), where recent negative trials were assumed to have high cognitive control demands, while novel negative critical trials had low demands.

#### 2.3.2 Imaging Analyses

All DICOM images were first converted to BIDS format (Gorgolewski et al., 2016) for each participant using in-house scripts (https://hub.docker.com/u/ccplabwustl) and subsequently preprocessed using FMRIPREP version 1.3.2 (Esteban et al., 2018; RRID: SCR_016216), a Nipype (Gorgolewski et al., 2011; RRID:SCR_002502) based tool. Each T1w (T1-weighted) volume was corrected for INU (intensity non-uniformity) using N4BiasFieldCorrection v2.1.0 (Tustison et al., 2010) and skull-stripped using antsBrainExtraction.sh v2.1.0 (using the OASIS template). Brain surfaces were reconstructed using recon-all from FreeSurfer v6.0.1 (RRID: SCR_001847), and the brain mask estimated previously was refined with a custom variation of the method to reconcile ANTs-derived and FreeSurfer-derived segmentations of the cortical gray-matter of Mindboggle (Klein et al., 2017; RRID: SCR_002438). Spatial normalization to the ICBM 152 Nonlinear Asymmetrical template version 2009c (RRID: SCR_008796) was performed through nonlinear registration with the antsRegistration tool of ANTs v2.1.0 (RRID: SCR_004757), using brain-extracted versions of both T1w volume and template. Brain tissue segmentation of cerebrospinal fluid (CSF), white-matter (WM) and gray-matter (GM) was performed on the brain-extracted T1w using fast (FSL v5.0.9, RRID: SCR_002823).

Functional data were slice time corrected using 3dTshift from AFNI v16.2.07 (RRID: SCR_005927) and motion corrected using mcflirt (FSL v5.0.9). Distortion correction was performed using 3dQwarp (AFNI v16.2.07). This was followed by co-registration to the corresponding T1w using boundary-based registration with six degrees of freedom, using bbregister (FreeSurfer v6.0.1). Motion correcting transformations, field distortion correcting warp, BOLD-to-T1w transformation and T1w-to-template (MNI) warp were concatenated and applied in a single step using antsApplyTransforms (ANTs v2.1.0) using Lanczos interpolation. Following FMRIPREP preprocessing, additional pre-processing occurred with AFNI software (Cox, 1996), including a frame-wise motion censoring (FD > 0.9 mm) and image normalization (i.e., demeaning).

Five participants failed behavioral quality control checks (e.g., more than 40% of trials and/or 5 trials in a row are not responded in one run of a task) and were excluded from analyses for the problematic task, that is, if participant failed quality control for one run of a task, both runs were excluded from analyses, as runs were concatenated in our analyses.

##### Univariate Analyses

The study utilized a mixed blocked / event-related design (Petersen & Dubis, 2012; Visscher et al., 2003), allowing for the separation of sustained (block-related) from transient event-related activation dynamics. After preprocessing, sustained and event-related effects were estimated using AFNI General Linear Model (GLM) functions (3dDeconvolve, 3dREMLfit) for each participant at the voxel-wise level (which includes 6-parameter motion and framewise censoring parameters, as well as polynomial detrending, using the -polort flag). The GLM estimation was conducted using a finite impulse response approach (Glover, 1999), implemented in AFNI using piecewise linear splines (“TENT” functions). This approach yielded an estimate of the full event-related time-course for each trial type. Timepoint estimates (“knots”) were set to 1 knot per 2 TRs. Voxel-wise beta coefficient estimates obtained from the GLM were assigned into cortical parcels based on the Schaefer 400 parcels, 7 networks atlas (Schaefer et al., 2018), and subcortical parcels via the CIFTI FreeSurfer segmentation (19 nuclei) (Glasser et al., 2013). Within both the high demand and high minus low (i.e., high-low) demand contrasts, we focused on the 2-TR “target” knot expected to show maximal high > low activation difference during each trial, representing the peak of cognitive control demand (Braver et al., 2020).

##### Definition of Parcels of Interest

We have previously demonstrated robust cross-task high-low control demand contrast in the DMCC dataset (Braver et al., 2020), the results of which we used to identify the two sets of brain parcels for the current analysis: first, a target set of 34, primarily located within the fronto-parietal regions (“FPN34”- 6762 voxels, Figure 2A), with significant greater activation in the high than the low control demand condition within each of the four tasks. Second, for a negative control, we identified a set of 34 parcels (“NULL34” – 6656 voxels, Figure 2B) without a significant activation difference between high and low control demand conditions (i.e., null effect) in Braver et al., 2020, setting an insignificance threshold of -1.5<t<1.5 in each of the four tasks. As expected, this set of 34 control parcels was located in regions not generally associated with cognitive control. A full list of these two parcel sets, with centroid coordinates, can be found in the supplementary materials (*ParcelsTable*).

**Figure 2.**
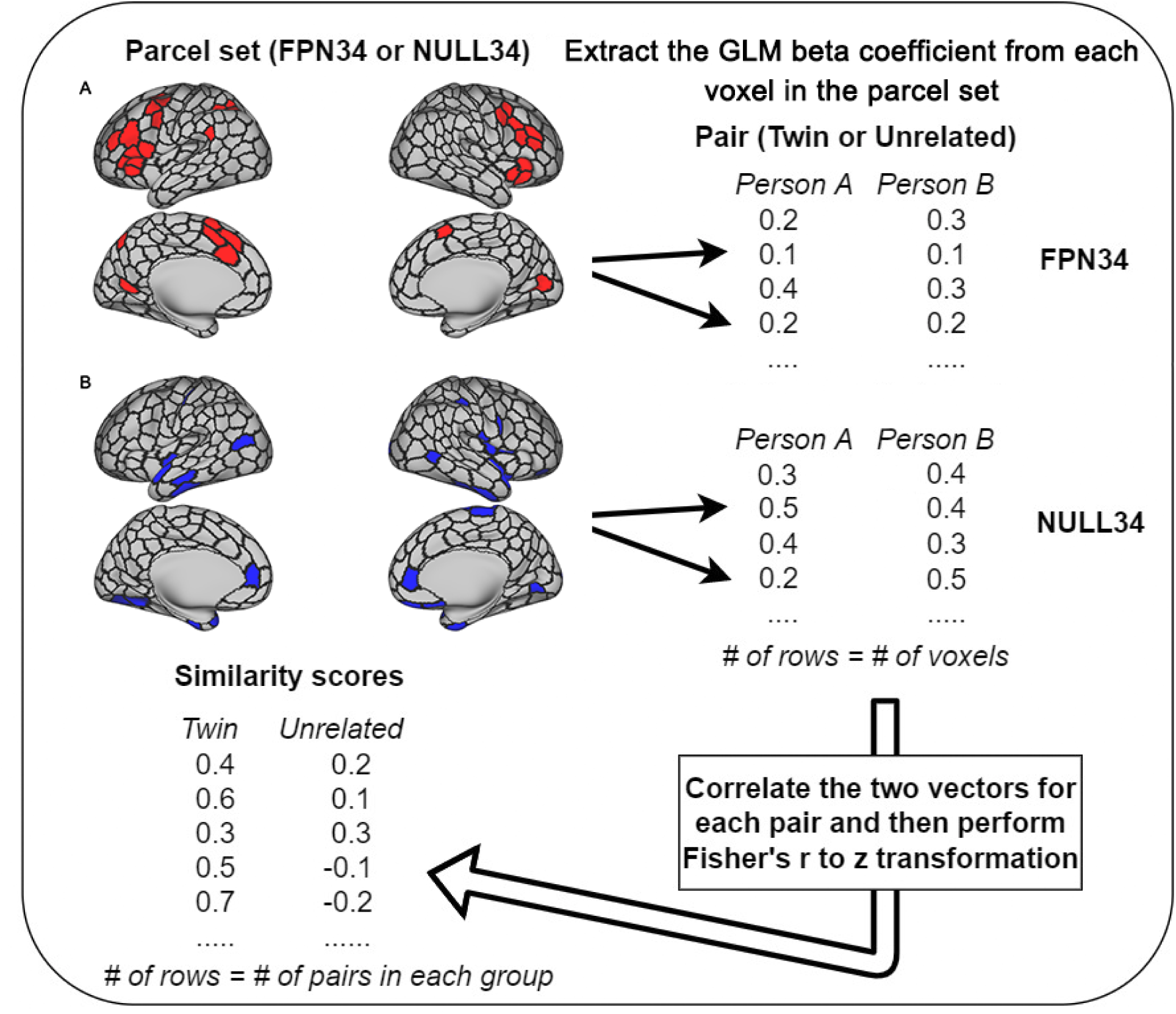
A Flowchart of the Twin Similarity Analyses - FPN34 (A) and NULL34 (B) Parcel Sets from Schaefer 400 Parcellation (visualized on a surface-based template with parcel boundaries emphasized)

##### Multivariate Analyses – Twin Similarity in Neural Activation Patterns

To compute twin similarity in activation patterns, we first extracted the beta coefficient estimates of all voxels obtained from univariate GLM analyses for each person, each parcel set (FPN34 or NULL34), and each contrast (high control trials or high - low control contrast), at the target knot of each task. Next, for each task, we performed Pearson’s correlations of vectors of these GLM beta coefficients within each pair (twin or unrelated), separately for each contrast and each parcel set. Consequently, two vectors of similarity scores (i.e., correlation coefficients) were generated – one for all twin pairs and the other for all unrelated pairs. Figure 2 is a visualization of these steps of analyses.

In addition to examining the twin similarity at the target knot, we are also interested in examining the temporal dynamics of twin similarity within task trials. Thus, we repeated the same steps shown in Figure 2, but at every 2-TR knot (i.e., timepoint) across task trials, not just the target knot. This allowed us to compare the activation patterns across the duration of a task trial to examine the potential temporal dynamics of pattern similarity. We performed these time-course analyses within each task separately, as they varied in duration (i.e., different number of TRs).

Lastly, to investigate the domain generality of the neural representations of cognitive control, we computed activation pattern similarity *between tasks* at the target knot. Specifically, rather than computing similarity scores for each participant pair (twin or unrelated) within each task, we performed Pearson’s correlations of vectors of GLM beta coefficients (from the target knot) between all possible task pairings (e.g., AX-CPT and Cued-TS, AX-CPT and Sternberg, Cued-TS and Sternberg) within each participant pair, separately for each contrast and each parcel set. We then averaged all pair-wise cross-task similarity scores to obtain a single similarity score representing cross-task pattern similarity for each participant pair.

##### Anatomic Similarity

To examine anatomic similarity within twin pairs and within unrelated pairs, we followed an approach first described in Polk et al (2007) and Pinel et al (2015). Specifically, we first extracted each person’s voxel-wise intensity values from the preprocessed anatomical T1-weighted scans for each parcel set and then computed anatomic similarity in each pair (twin or unrelated), separately for each parcel set, through Pearson’s correlations of voxel-wise intensity vectors. Consequently, we obtained two sets of anatomic similarity scores (i.e., correlation coefficients) – one for all twin pairs and the other for all unrelated pairs.

#### 2.3.3 Statistical Analyses

A summary of primary statistical analyses can be found in the supplementary materials (*StatisticalAnalysesFlowchart*). All analyses were performed using R version 3.6.3 (R Development Core Team 2015). Robust t-tests (Yuen, 1974) were conducted using the *DescTools* package (https://cran.r-project.org/web/packages/DescTools/index.html), with trimmed means (0.1 trim) to reduce the effects of outliers and non-normality. Linear mixed models were performed using the *nlme* package (https://cran.r-project.org/web/packages/nlme/index.html). Post-hoc contrasts of linear mixed models were conducted using the *lsmeans* package (https://cran.r-project.org/web/packages/lsmeans/index.html). The Fisher r-to-z transformation was applied to all correlation coefficients. Code for replicating the figures and analyses is available at the Open Science Foundation, https://osf.io/9d7ym/.

## 3. Results

### 3.1 Behavioral Task Performance

We behaviorally validated that our trial type contrasts indexed cognitive control demands. Specifically, we tested the assumption that cognitive control demand would be associated with increased behavioral interference, observed as a decline in task performance, i.e., slower RTs and higher error rates present on high demand relative to low demand trials. To test for control demand effects, for each task we conducted paired Yuen robust t-tests (two-tailed) on both error rates and RT (calculated from accurate trials only) in twins and unrelated individuals (i.e. does not include twins) separately.

As expected, we found highly robust interference effects (Table 1) in all four tasks, such that RT and error rates were higher on trials with high than low control demands. All effects were highly significant in twins: Stroop (RT: t = 14.0, p<0.0001; error: t = 3.2, p=0.0048), AX-CPT (RT: t = 6.6, p<0.0001; error rate: t = 5.7, p<0.0001), Cued-TS (RT: t = 5.7, p<0.0001; error: t = 4.9, p<0.0001), Sternberg (RT: t = 7.3, p<0.0001; error rate: t = 11.2, p<0.0001). Similarly, all effects were significant in unrelated individuals: Stroop (RT: t = 12.9, p<0.0001; error: t = 3.8, p=0.0007), AX-CPT (RT: t = 7.5, p<0.0001; error rate: t = 4.9, p<0.0001), Cued-TS (RT: t = 2.2, p=0.0317; error: t = 4.9, p<0.0001), Sternberg (RT: t = 4.9, p<0.0001; error rate: t = 5.9, p<0.0001). Together, these results support using these selected trial types to index high control and low control demand in each task.

**Table 1:**
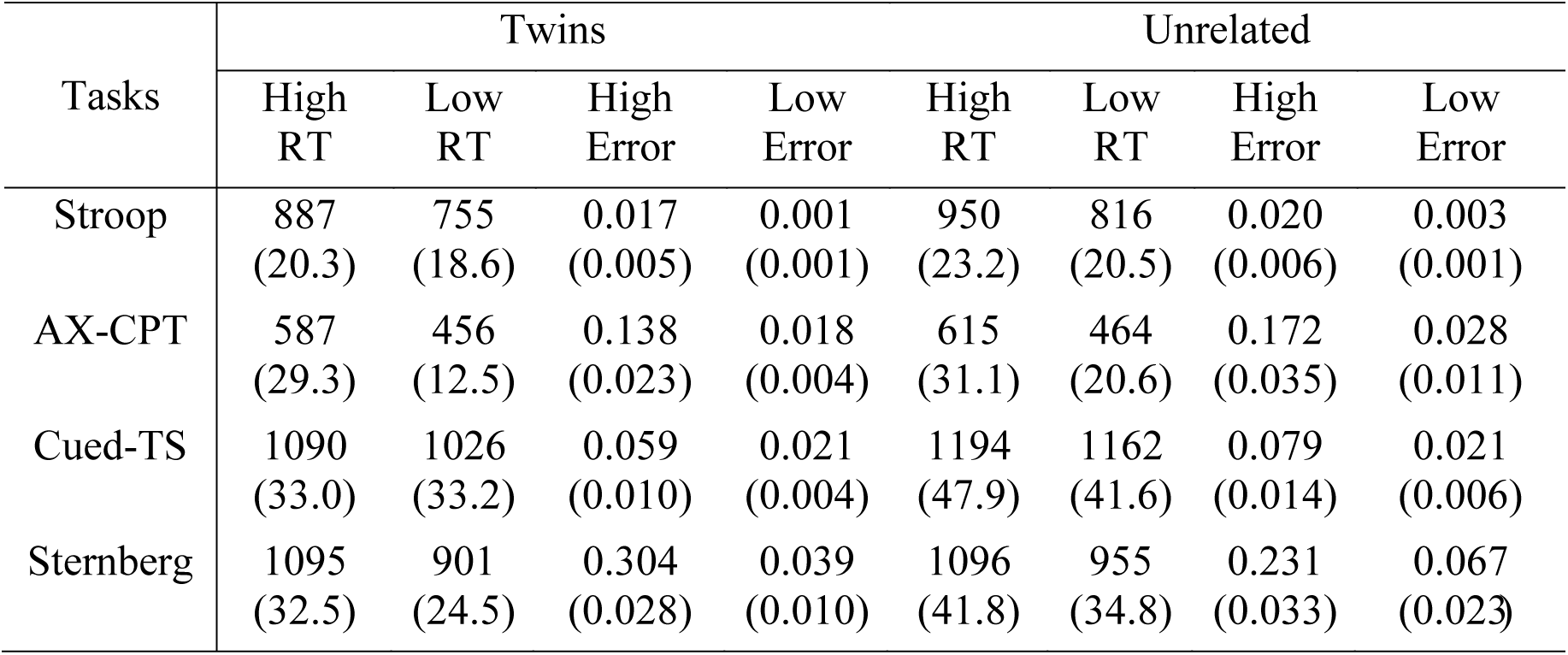
Robust Mean (Standard Error of the Mean) of Task Performance for High and Low Demand Trials in Twins and Unrelated Individuals

We also compared the interference effects between twins and unrelated individuals by conducting two-sample robust t-tests (two-tailed) on RT and error rates. No significant difference in interference effects was detected in RT or error rates (p>0.05), except that the interference effect on error rates in Sternberg was greater in twins than in unrelated individuals (t = 2.6, p=0.0116). Together, the behavioral data suggest that twins and unrelated individuals responded to cognitive control demands in a mostly equivalent manner.

### 3.2 Univariate Analyses: Timecourses of Neural Activations

As a first validation step, beta coefficients from univariate GLM analyses were extracted, and plotted as event-related timecourses for each group. Based on visual inspection of the timecourses of neural activation for both high control demand trials and high-low control contrast in twins and unrelated pairs (see Figure 3), it is clear that the two groups had essentially equivalent neural activation in each task across all timepoints. To confirm this pattern, we conducted robust t-tests on the mean beta estimates of the two groups; as expected, there was no significant difference in activation between the two pair groups at any timepoint in each of the four tasks (p>0.05). Thus, any observed differences between twin and unrelated pairs is unlikely to be explained by overall activation differences related to cognitive control demand among the two participant groups.

**Figure 3.**
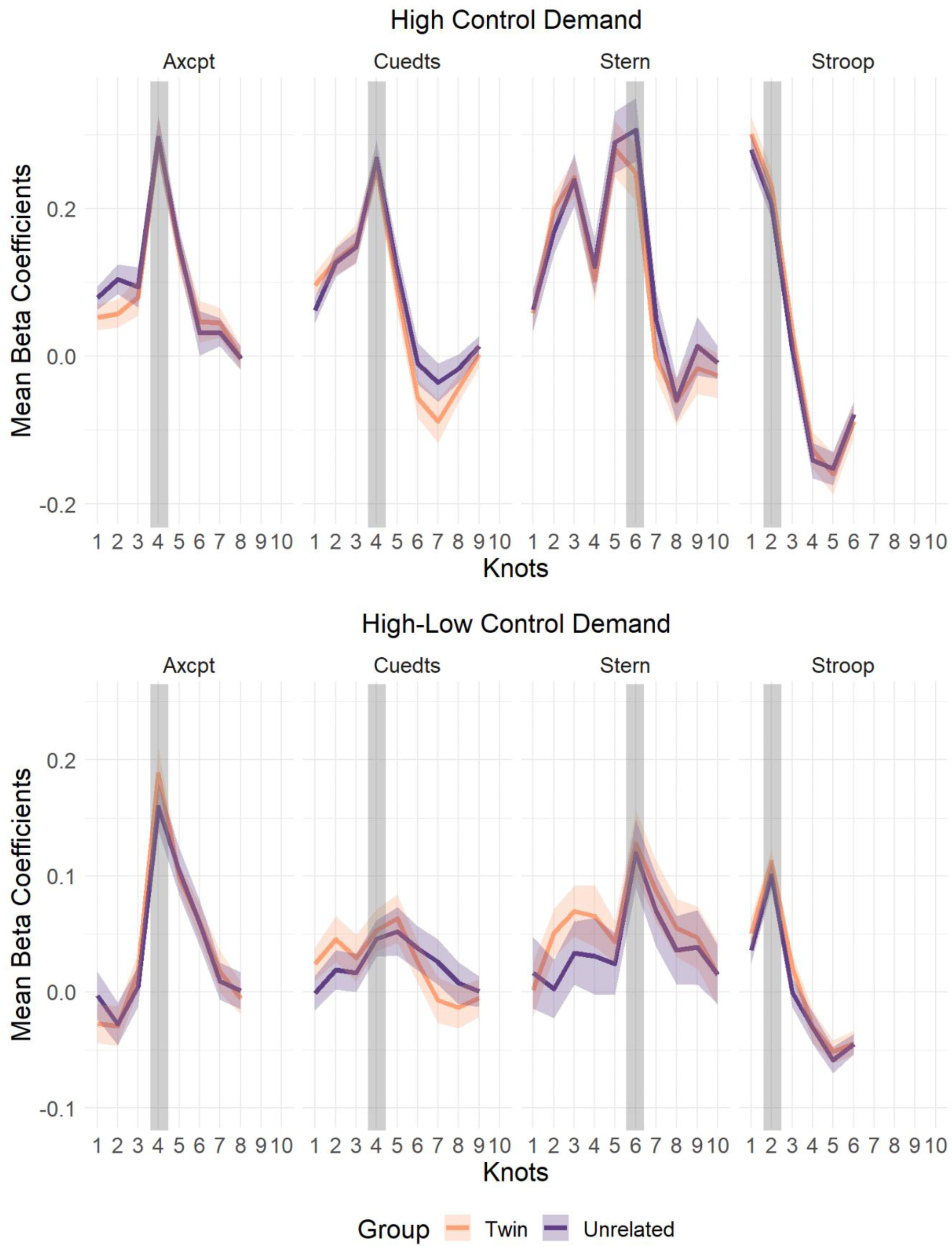
Robust Mean and Standard Error of the Mean of Univariate Beta Timecourse Coefficients of Twins and Unrelated Pairs by Task

### 3.3 Multivariate Pattern Similarity Analyses: Overview

The prior analyses focused on univariate activity. The remaining analyses use *pattern similarity* approaches to contrast twin and unrelated pairs. A summary of the statistical analyses can be found in the supplementary materials (*StatisticalAnalysesFlowchart*). The first set of analyses focused on high control demand trials, whereas the second set focused on the high – low control demand contrast to provide greater specificity. Within each set, four inter-related analyses were conducted which examined twin effects at progressively greater levels of detail. First, we aggregated the similarity scores across all four tasks and investigated the overall effects of twinness (i.e., twins and unrelated) and parcel set (i.e., FPN34 and NULL34) on patterns of activation. Second, we compared similarity scores within each of the four tasks individually. Third, we examined the temporal dynamics of pattern similarity for each of the four tasks, specifically the activation pattern similarity at each timepoint (2-TR knot) within the trials. Fourth, we investigated the effect of twinness on cross-task similarity scores to test the hypothesis that twin similarity effects are domain-general with respect to cognitive control demand. Finally, we conducted a control analysis to compare task activation pattern similarity with *anatomic* similarity in twins and unrelated pairs.

#### 3.3.1 Twin vs. Unrelated Similarity in High Control Demand Trials

In the first set of pattern similarity analyses, we took advantage of the multi-task event-related design to examine neural activation patterns selectively on high control demand trials. We predicted that would observe increased activation pattern similarity in twins relative unrelated pairs, but that this effect would be anatomically specific, and so reliably stronger in the FPN34 set than in the NULL34 set.

##### Linear Mixed Model: Aggregating across Tasks

Aggregating the similarity scores of all four tasks, we built a linear mixed model to test for the fixed effects of twinness (i.e., whether the pair is twin or unrelated) and parcel set (FPN34 or NULL34), as well as their interaction, on pattern similarity in high control demand trials. We also included task (AX-CPT, Cued-TS, Sternberg, and Stroop) and pair ID (all twin and unrelated pairs) as random effects in the model, to account for potential variance due to each individual pair and task. The detailed model set up can be found in the supplementary materials (*HighControl*).

We identified a highly significant parcel set by twinness interaction (p<0.0001), which we further investigated with post-hoc contrasts. Specifically, we hypothesized that the interaction was due to a significant difference in similarity between the twin and unrelated pairs in the FPN34 parcel set, but not in the NULL34 parcel set, as NULL34 is not functionally relevant for cognitive control and should only exhibit arbitrary patterns of similarity within each pair. This hypothesis was mostly confirmed (see Figure 4), in that the similarity differences were highly significant in the FPN34 (p_corrected_<0.0001). In particular, not only were overall similarity scores higher in the FPN34 parcel set relative to the NULL34 parcel set, but also the effect of twinness was significantly greater. However, in the NULL34 parcel, the effect of twinness, though much weaker than FPN34, was still marginally significant (p_corrected_=0.0463). Nevertheless, the overall pattern suggests that the twin effect on pattern similarity in high control demand trials was primarily expressed in the FPN34 parcel set.

**Figure 4:**
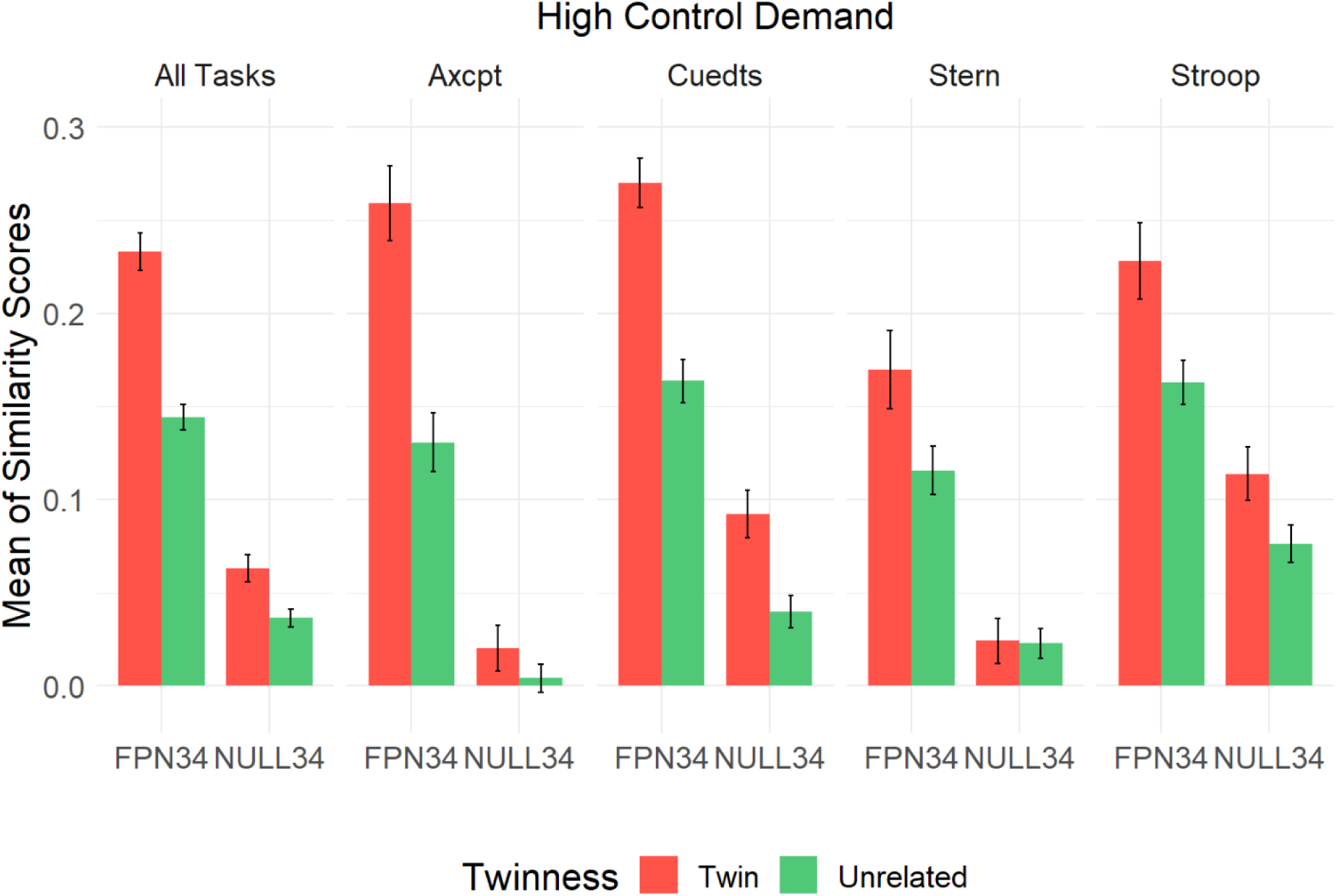
Mean (and Standard Error of the Mean) of Similarity Scores in High Control Demand Trials

##### Linear Mixed Model: Within Each Task

We next examined the similarity of twins and unrelated individuals within each task separately, again using linear mixed models. The model and output for each task can be found in the supplementary materials (*HighControlByTask*). In three of the four tasks (not in Stroop), the interaction of twinness and parcel set was statistically significant, so we carried out post-hoc analyses within each of the three tasks. Consistent with the trends we observed above, twins had significantly higher FPN34 parcel set activation pattern similarity than unrelated individuals in the (see Figure 4) in AX-CPT (p_corrected_<0.0001), Cued-TS (p_corrected_<0.0001), and Sternberg (p_corrected_=0.0219). Conversely, NULL34 activation pattern similarity did not differ significantly in AX-CPT or Sternberg. However, for Cued-TS, a significant difference was observed in the NULL34 parcel set (p_corrected_=0.0087). For Stroop, although there was no significant interaction, to examine the direction of the similarity difference between twins and unrelated pairs, we built linear mixed models for FPN34 and NULL34 similarity scores separately, with twinness (twins or unrelated) as the only fixed effect and pair ID as the only random effect. Like in the other tasks, we found significantly greater twin similarity relative to unrelated pairs in both FPN34 (p=0.0043) and NULL34 (p=0.0328) in Stroop. Overall, the results support the hypothesis that the twin effect was largely specific to the FPN34 parcel set, and absent or weaker in the NULL34 parcel set when examined separately in each task.

##### Two-sample T-tests: Temporal Dynamics of Twin vs. Unrelated Similarity

Next, we conducted robust two-sample t-tests (two-tailed) to compare similarity scores of twins and unrelated pairs at every timepoint estimate (2-TR knot) within each of the four tasks. The t-scores indicate the degree to which the neural pattern similarity of twins differs from that of unrelated pairs, with higher t scores indicating greater differences and higher twin similarity. Here, linear mixed models were not carried out as we were interested in similarity differences between twins and unrelated for each parcel set individually, rather than the parcel set by twinness interaction. We plotted these t-scores across the task trials to illustrate the temporal dynamics of the twin similarity effect (see Figure 5), with the target timepoint (expected peak control demand) shaded in grey and a red line illustrating the statistical significance threshold (p<0.05). In all four tasks, the similarity scores at the target timepoint in these high control demand trials were significantly higher in twins than in unrelated pairs in the FPN34 parcel set. Notably, in AX-CPT and Cued-TS, the twin effect (i.e., twin > unrelated) peaked at the target timepoint relative to the other timepoints in the FPN34 parcel set. In Sternberg and Stroop, the twin effect was strong in the FPN34 parcel set but did not peak at the target timepoint. Finally, except for Cued-TS, in which there was a peak at the target timepoint in the NULL34 parcel set, the NULL34 twin effect in all tasks was rather weak and non-specific in terms of peak timepoint. Likewise, and consistent with the prior analyses, the twin effect was weaker in NULL34 than FPN34 across most timepoints.

**Figure 5:**
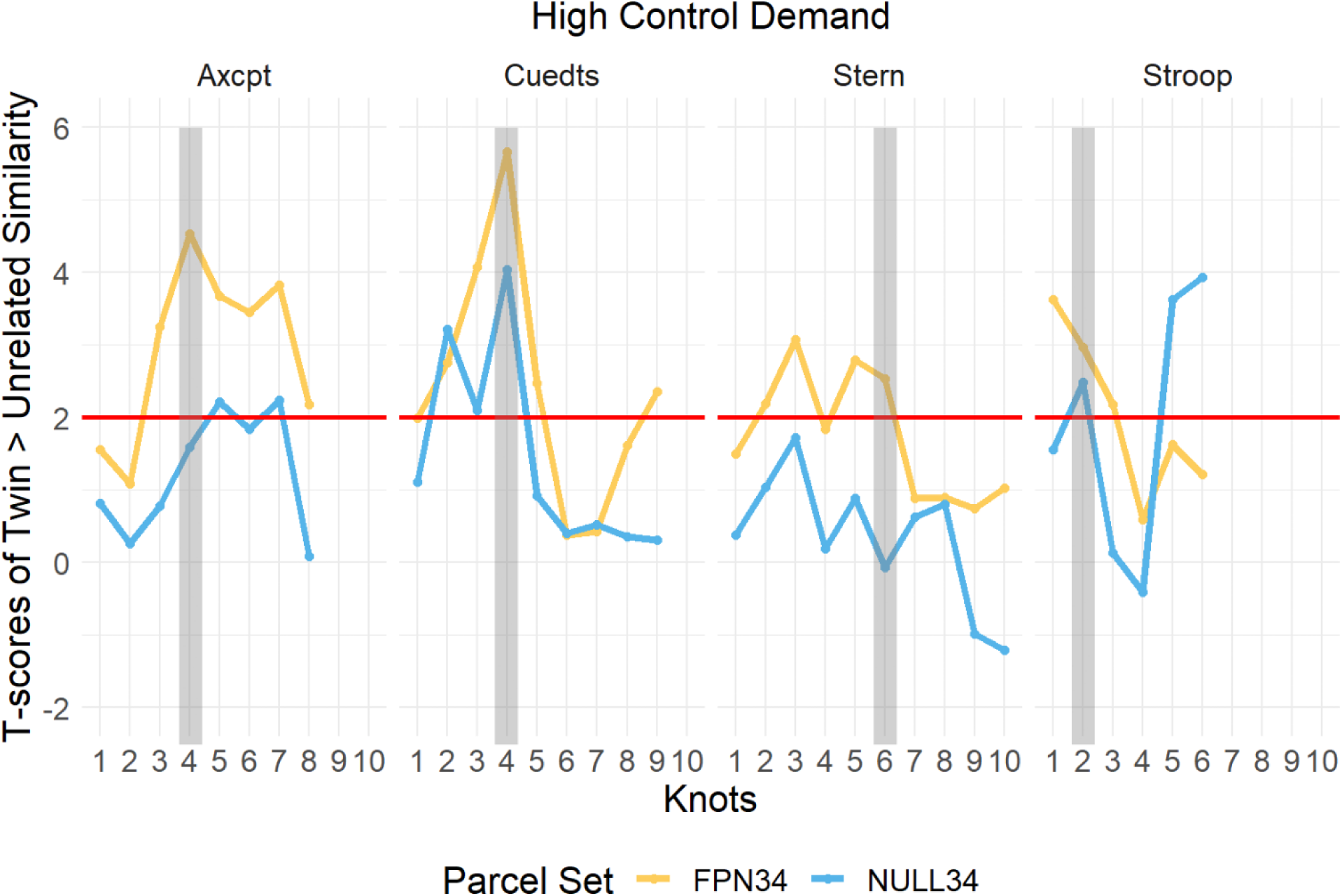
Temporal Dynamics of Difference in Pattern Similarity (Twin > Unrelated) in High Control Demand Trials

##### Linear Mixed Model: Cross-task Twin Similarity

As an alternative approach to examine the domain generality of cognitive control, we computed cross-task similarity in neural activation patterns for twins and unrelated pairs, averaging all pair-wise comparisons among the four tasks to obtain a cross-task similarity measure for each pair. We then built the same linear mixed model as described above to investigate whether twins exhibited greater activation pattern similarity than unrelated pairs in the FPN34 parcel set but not in the NULL34 parcel set. The detailed model set up can be found in the supplementary materials (*HighControlCrossTask*). Consistent with the trends observed in within-task pattern similarity (see Figure 6), a significant parcel set by twinness interaction (p<0.0034) was detected. Moreover, post-hoc tests indicated that the similarity differences between twins and unrelated pairs were highly significant in the FPN34 (p_corrected_<0.0001), but not in the NULL34 (p_corrected_<0.2085), suggesting robust twin effects on pattern similarity even across tasks, which were again specific to the FPN34 parcel.

**Figure 6:**
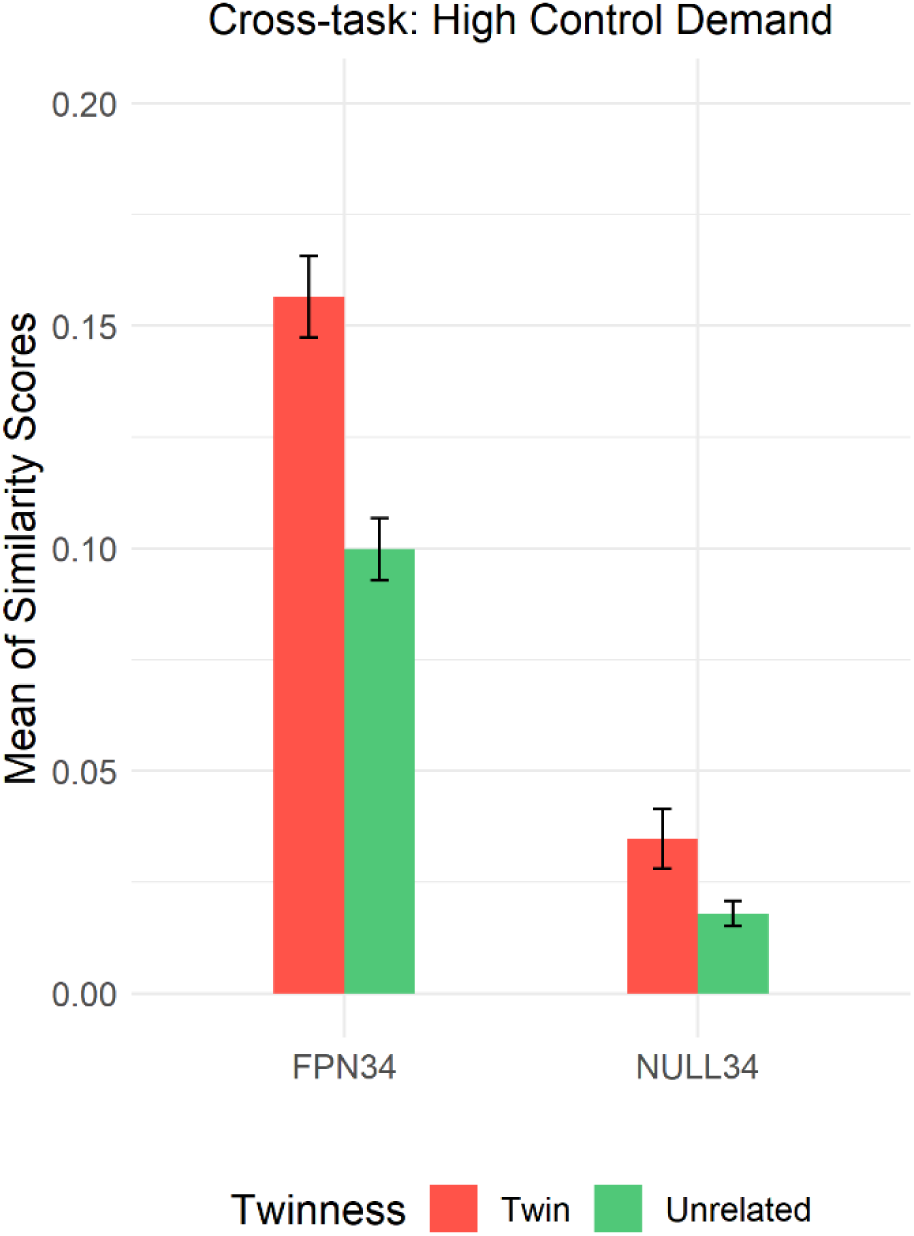
Mean (and Standard Error of the Mean) of Cross-task Similarity Scores in High Control Demand Trials

#### 3.3.2 Twin vs. Unrelated Similarity in High-Low Control Demand Contrast

In the second set of multivariate analyses, we examined neural activation patterns in terms of the high-low control demand contrast, a more stringent examination of the twin similarity effect. Unlike the above analyses that focused only on high control demand trials, here we tested whether the differences in neural activation patterns between high and low control demand trials were also similar in twins and specific to the FPN34 parcel set. It should be noted that this statistical analysis approach will inevitably increase the amount of measurement error and also impose stricter constraints (i.e., looking for similarity in activation difference patterns). Thus, purely on a priori statistical grounds, this set of analyses should yield less robust results relative to those found when examining high control demand trials alone. Nevertheless, since this analysis more strongly isolates the effect of cognitive control demand itself (by removing activation variation due to general engagement on task trials), it provides a stronger test of our hypotheses.

##### Linear Mixed Model: Aggregating across Tasks

Similar to the analyses performed in high control demand trials, we built a linear mixed model to examine the effects of twinness and parcel set when aggregating the four tasks, as well as their interaction effect on the similarity of activation patterns in the high-low control demand contrast. The detailed model set up can be found in the supplementary materials (*High-Low Control*). Consistent with our hypothesis, the results paralleled those we found with high control demand trials, with a significant twinness by parcel set interaction (p=0.0023), which we followed up with post-hoc contrasts.

As predicted, we again observed a differential twin effect on the similarity scores in the two parcels. Importantly, the difference in similarity scores between twins and unrelated individuals in the high-low control contrast was significant for the FPN34 parcel set (p_corrected_<0.0001) but not even close to significant for the NULL34 parcel set (p_corrected_=0.9364). These results indicate that twins still exhibited greater pattern similarity than unrelated pairs, even when considering the neural activation difference between high and low control demand trials (see Figure 7).

**Figure 7:**
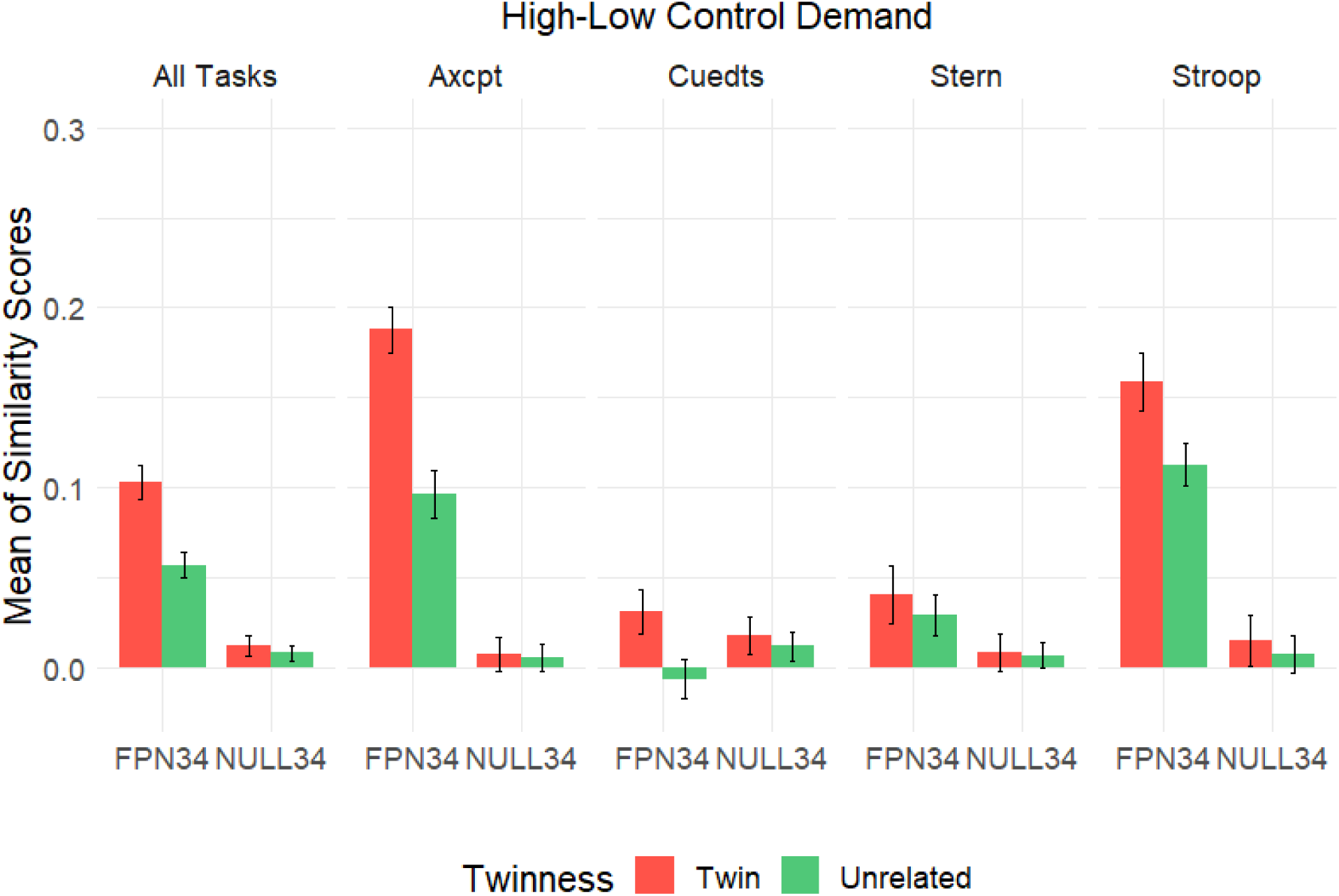
Mean (and Standard Error of the Mean) of Similarity Scores in High-Low Control Demand Contrast

##### Linear Mixed Model: Within Each Task

Next, we computed the differences in pattern similarity within individual tasks (see Figure 7). The models for each task can be found in the supplementary materials (*High-LowControlByTask*). We detected a significant twinness by parcel set interaction in AX-CPT, and followed up with post-hoc contrasts. In AX-CPT, twins had significantly higher similarity in activation patterns than unrelated pairs in the FPN34 parcel set (p_corrected_<0.0001) but not in the NULL34 parcel set (p_corrected_=0.9987). For the other three tasks, there was no significant interaction. However, similar to what we did above, we built two linear mixed models for FPN34 and NULL34 in each of the three tasks to examine the direction of our hypothesis, that is, if twins were higher in pattern similarity than unrelated pairs in the FPN34 parcel set, but not the NULL34 parcel set. This hypothesis was again confirmed, in that we were able to detect significant differences in similarity scores for Cued-TS (p=0.0292) and Stroop (p=0.0245) in the FPN34 parcel set, but not in the NULL34 parcel set. For Sternberg, although the difference in pattern similarity was not significant for either of the two parcel sets, the general profile of the data was the same, with twins numerically higher than unrelated individuals in similarity scores, and the effect numerically larger in FPN34 than NULL34. Finally, because the magnitude of similarity scores was small in Cued-TS and Sternberg relative to the other two tasks, we conducted one-sample t-tests to confirm that the similarity scores of twins were significantly greater than zero for FPN34 in Cued-TS (p=0.009) and Sternberg (p=0.0089). Overall, even at the level of individual tasks, the twins exhibited greater similarity in activation patterns than unrelated individuals under increased cognitive control demand, an effect largely specific to the FPN34 parcel set.

##### Two-sample T-tests: Temporal Dynamics of Twin Similarity

We performed a robust two-sample t-test (two-tailed) to compare similarity scores of twins and unrelated pairs in the high-low control demand contrast at every timepoint. Again, even with this more stringent test, the difference in similarity scores between twins and unrelated pairs at the target timepoint were significant in three of the four tasks (not Sternberg). Critically, the twin effect also peaked at the target timepoint in the FPN34 parcel set for these same three tasks; whereas this temporal profile was not observed in the NULL34 parcel set (see Figure 8). Moreover, in the NULL34 parcel set, there was no significant effect of twinness (i.e. twins > unrelated) at any timepoint.

**Figure 8:**
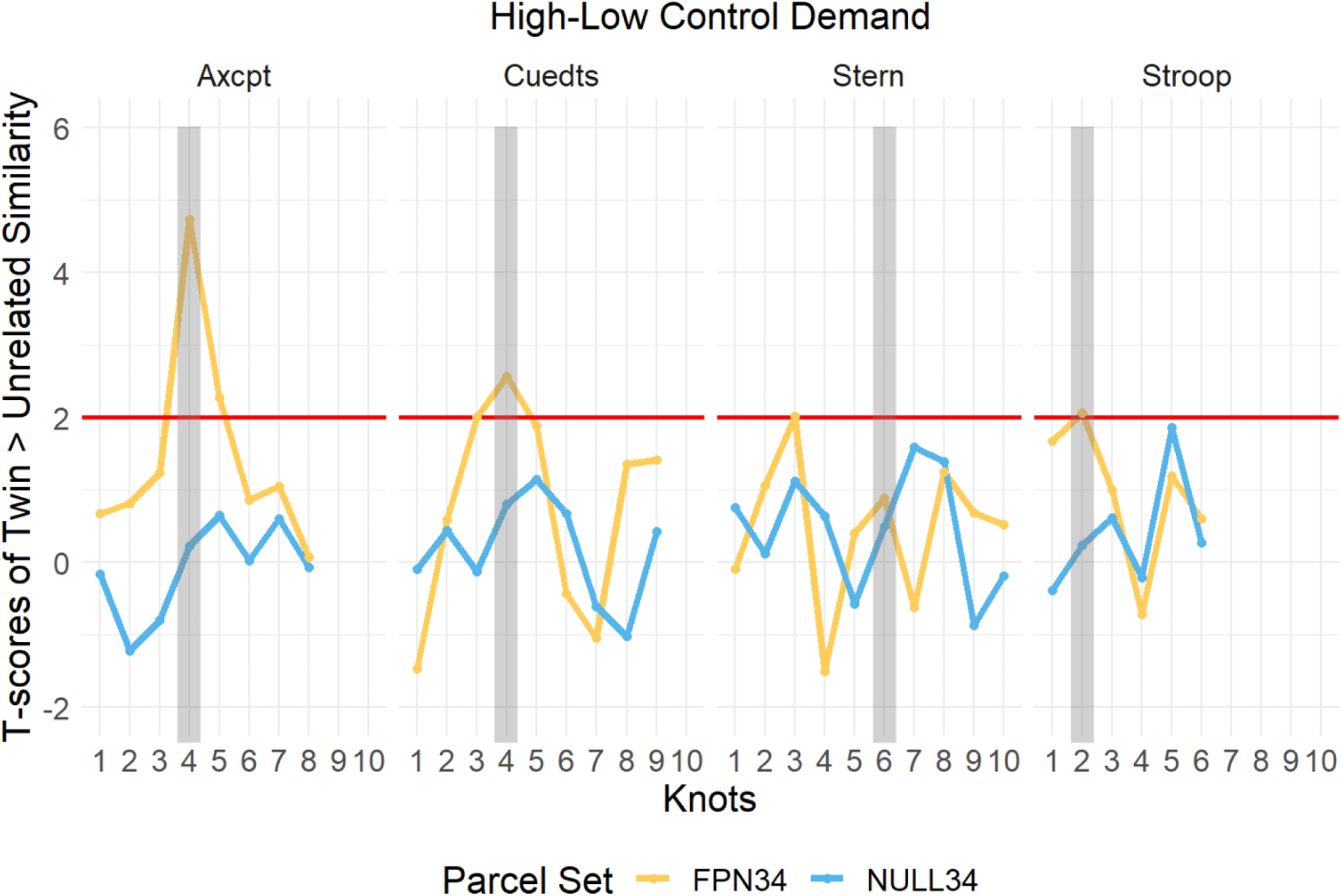
Temporal Dynamics of Difference in Pattern Similarity (Twin > Unrelated) in High-Low Control Demand Contrast

A potential concern of comparing the NULL34 set with FPN34 is that we hand-selected the NULL34 parcels based on t-values from univariate analyses. Consequently, there could be potential biases introduced from the use of these parcels due to non-specific factors (e.g., low SNR). Therefore, we also ran the analyses with parcels from the somatomotor network (10819 voxels) as the control parcel set, since these are anatomically defined and pre-specified, and so unbiased with respect to our hypothesis; details of these results are presented in supplementary materials. The findings were very similar with this control parcel set: a significant twinness by parcel set interaction was observed in both high control trials (*HighControlSOMA*) and high-low demand contrast (*High-LowControlSOMA)* at the target timepoint. For time-course analyses, the pattern similarity of the somatomotor network was very much like that of NULL34, in that it did not demonstrate specificity to increased control demand in any of the four tasks.

##### Linear Mixed Model: Cross-task Twin Similarity

The fourth set of analyses examined the effect of twinness on cross-task similarity scores, by means of linear mixed models. The model can be found in the supplementary materials (*High-LowControlCrossTask*). Although there was no significant parcel by twinness interaction, we built two linear mixed models (one each for FPN34 and NULL34) to examine the direction of our hypothesis; that is, if twins were higher in cross-task pattern similarity than unrelated pairs in the FPN34 parcel set, but not the NULL34 parcel set. This hypothesis was again confirmed, in that we were able to detect significant differences in similarity scores in the FPN34 (p=0.023) but not in the NULL34 parcel set (p=0.6077) (see Figure 9). Additionally, we ran the cross-task pattern similarity analyses with parcels from the somatomotor network as another control parcel set. The findings were again very similar with the NULL34 control parcel set in both high control trials (*HighControlSOMA*) and high-low demand contrast (*High-LowControlSOMA)* at the target timepoint. Overall, these results suggested that the twin effects were pronounced even in cross-task activation pattern similarity during increased cognitive control. Moreover, the effects were specific to the FPN34 parcel set, a trend that is consistent with what was observed in within-task pattern similarity.

**Figure 9:**
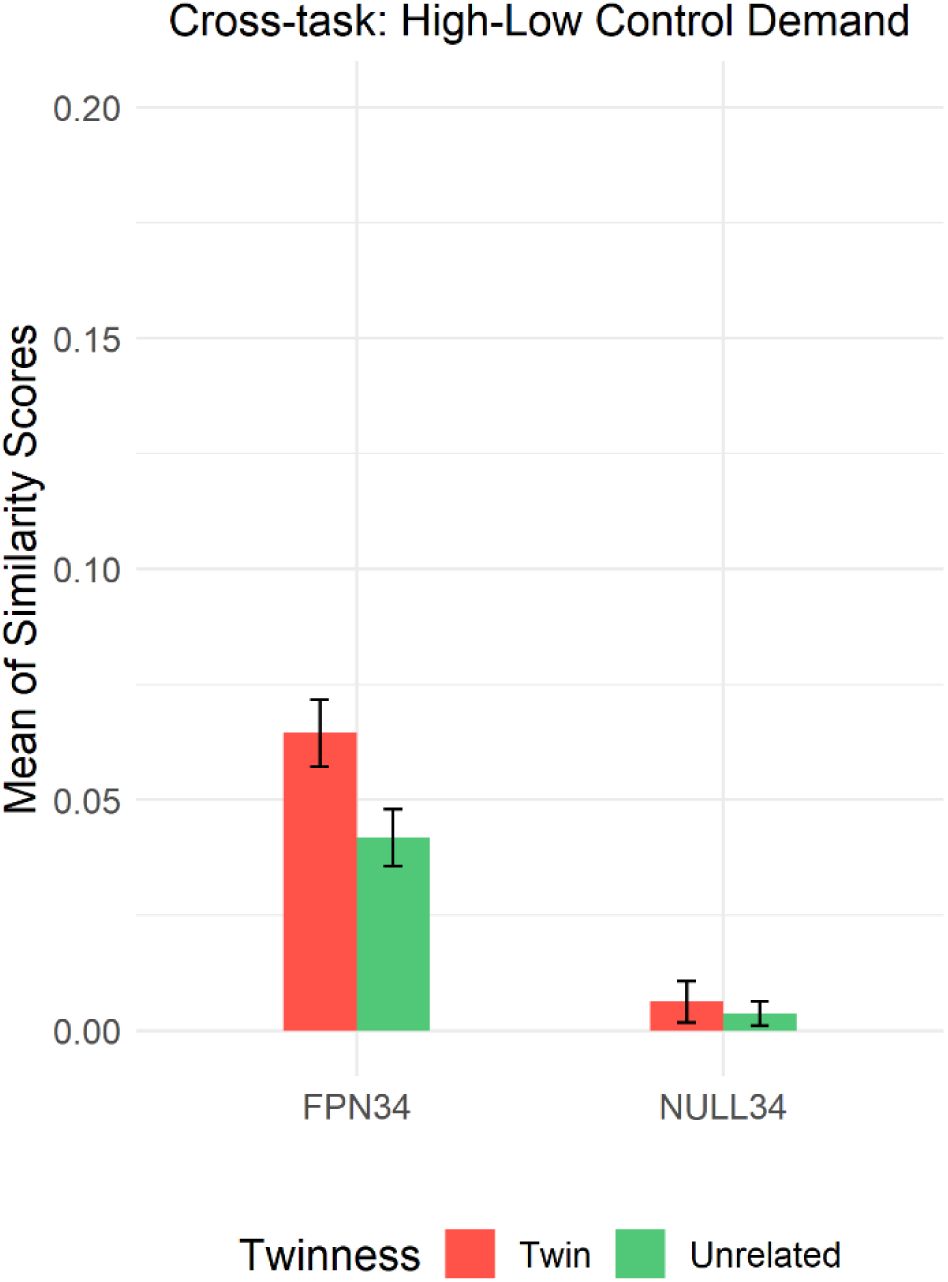
Mean (and Standard Error of the Mean) of Cross-task Similarity High-Low Control Demand Contrast

**Figure 10:**
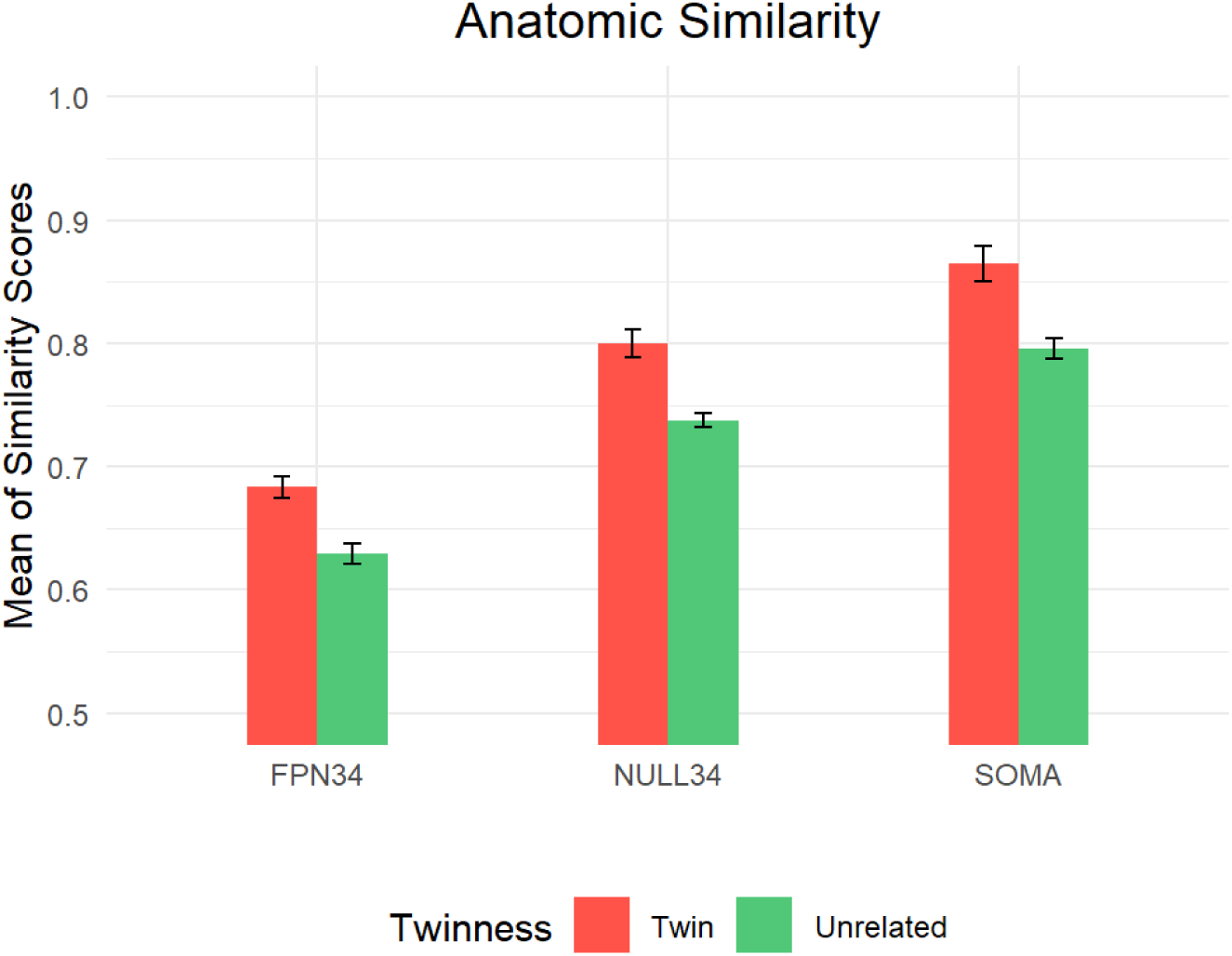
Anatomic Similarity (z-transformed) in Twins and Unrelated Pairs by Parcel Set.

#### 3.3.3 The Relationship of Functional to Anatomic Similarity

Finally, we conducted an analysis to explore the influence of anatomic similarity, given that MZ twin pairs have more similar brains than do unrelated people. A potential confound in the observed twin effects in functional activation pattern similarity is that these effects could be potentially due to anatomic similarity within the regions of interest among monozygotic twin pairs. In particular, it is possible that the preferential effects that were observed in FPN34 might be due to increased anatomic similarity among twin pairs in this set of regions. To address this question, we followed an approach used in the prior literature (Pinel et al., 2015; Polk et al., 2007), in which the voxel-wise intensity patterns in anatomical scans were compared across pairs to determine anatomic similarity. Consequently, we utilized the T1-weighted scans to compute the anatomic similarity in voxel-wise intensity separately for twins and unrelated pairs.

Consistent with prior findings (Pinel et al., 2015; Polk et al., 2007), we were able to detect a consistent pattern of higher anatomic similarity among twins relative to unrelated pairs in each of the three parcel sets using repeated-measures ANOVA (main effect of twinness: *F*(1,73) = 38.31, p <0.0001; twin effect in each parcel set separately was significant at p < .001; see Figure 8). Moreover, the findings also indicated that there was no preferential effect in FPN34 over NULL34 and somatomotor network parcels (twinness x parcel set interaction, F(2,146) < 1); in fact, anatomic similarity was the lowest in FPN34 (main effect of parcel type: *F*(2,146) = 283.23, p <0.0001). This pattern counters the idea that the FPN34 effects observed in activation pattern similarity might reflect preferential anatomic similarity in this region. As a final analysis, we computed the relationship between anatomic and functional similarity across all pairs, both twin and unrelated, and all three parcel sets. In none of these was a significant positive correlation observed (all r’s < 0.37). Details of these correlational results and plots found in the supplementary materials (*AnatomicalCorrelations*). Taken together, these results provide additional evidence that the anatomic similarity among twins is unlikely to be the underlying source of variation that accounts for the twin similarity effect in cognitive control-related neural activation patterns, particularly for the FPN34.

## 4. Discussion

Understanding individual differences in cognitive control function is critical for both basic and translational research. The present work builds on prior work using multi-task, twin-based designs to examine cognitive control (Friedman et al., 2008; Miyake & Friedman, 2012), by extending these from behavioral analyses into the neuroimaging domain. In particular, we utilized a distinct and theoretically-focused cognitive control task battery to both conceptually replicate prior findings, but also demonstrate how novel neural measures can be harnessed to provide new evidence supporting a genetic contribution to individual differences in the brain networks that support cognitive control function. The results provide the first evidence that twin similarity in the neural representation of cognitive control is domain-general, but also exhibits both anatomic specificity to fronto-parietal regions and temporal specificity to the time of highest control demand within task trials. Together, our findings suggest that the fMRI activation patterns indexing cognitive control can be considered akin to a ‘neural fingerprint’, which reflects the fine-grained systemic, yet idiosyncratic variation in neural activity within fronto-parietal regions that characterize individuals, and their genetically-identical twins. More generally, our work highlights the potential of twin-based neuroimaging investigations and multivariate pattern analysis approaches for exploring heritability questions within this domain.

### 4.1 Domain-generality of twin similarity

Current theoretical and empirical work has suggested a domain-general cognitive control capacity (Assem et al., 2020; Kan et al., 2013; Miyake & Friedman, 2012), which enables flexible adaptation of thoughts and actions to fulfill a variety of task goals and demands (Cole et al., 2013; Braver, 2012). Yet this hypothesis could not be tested in prior studies using twin-based neuroimaging designs, since they relied on evaluating heritability of neural activation in single cognitive control tasks (Etzel et al., 2020; Koten et al., 2009; Matthews et al., 2007). Our work significantly extends these prior findings by demonstrating that twin similarity effects generalize across multiple cognitive control tasks; tasks which tap into different processes within this domain (e.g., selective attention, task-switching, working memory). In particular, we showed that not only did twins exhibit greater similarity within each of the four tasks, but they also had more similar cross-task activation patterns than did unrelated pairs of individuals, suggesting that there may be a domain-general yet idiosyncratic neural activation profile common to cognitive control processes, which is at least partly influenced by genetics.

In this regard, our neuroimaging design parallels that taken in prior behavioral studies, conducted in both adolescents (Friedman et al., 2008) and young adults (Friedman et al., 2016), in which multiple tasks tapping into cognitive control or executive functions are utilized (Friedman et al., 2008), and in more recent neuroimaging work from this same group, adding a focus on resting-state connectivity (Reineberg et al., 2018; 2020; Menardi et al., 2021). Yet the current study also departs from this prior work in a number of critical methodological aspects. In particular, Friedman and colleagues utilized latent variable methods to identify variation that is common to multiple tasks. Although latent variable approaches are well-established, they can be somewhat prohibitive for neuroimaging studies, due to their large-sample size constraints (Cooper et al., 2019). Thus, they will be an important direction for future work, in which large-scale twin neuroimaging and multi-task designs can be employed. In the current work, because of the smaller sample size, we chose to focus on multivariate cognitive control activation patterns, due to the flexibility and power of such approaches for addressing questions related to the neural coding and domain-generality of cognitive control (Freund, Etzel, & Braver, 2021). In particular, these neural activation pattern similarity methods are quite amenable to extensions that directly incorporate both within-task and cross-task similarity, and are also highly sensitive in capturing individual variation in neural activation patterns.

### 4.2 Anatomic specificity of twin similarity

Neuroimaging studies have consistently identified the frontal and parietal regions as involved in a wide range of cognitive control tasks, including but not limited to working memory, task-switching, and conflict resolution (Cole & Schneider, 2007; Cole et al., 2013; Dosenbach et al., 2006; Fedorenko et al., 2013; Hugdahl et al., 2015). Although most of this work comes from univariate activation and functional connectivity studies, a recent meta-analysis of MVPA studies indicated that the neural activation patterns within these regions are highly relevant for the encoding and representation of various task-relevant information (Woolgar et al., 2016). Our findings confirmed these fronto-parietal regions as a domain-general cognitive “core” (Duncan, 2010; Assem et al., 2020) engaged by cognitive control demands in all four tasks. More importantly, we showed that twin similarity in neural activation patterns was specific to the FPN parcel set (including regions overlapped with the multiple-demand system shown in Assem et al., 2020), while being unreliable in the set of parcels selected based on an absence of cognitive control effects (from univariate analyses). It is important to note that although it could be argued that the NULL parcel set was unfairly biased in favor of our hypothesis due to this preselection, we also found similar specificity to the FPN when we compared to a somatomotor network that had been pre-defined based on anatomy and connectivity, rather than due to its activation profile in our cognitive control tasks. Moreover, we found that although all three sets of parcels also exhibited higher *anatomic similarity* among twins than unrelated pairs, this was not preferential to the FPN parcels; in fact, in the FPN anatomic similarity was the lowest among all pairs, suggesting that increased levels of functional similarity in neural activation patterns among twins is simply not a byproduct of their anatomic similarity.

Our results also support prior work suggesting that fronto-parietal regions are a key locus of individual and genetic variation. Individual differences in higher cognitive processes and general fluid intelligence have been closely linked to variation in the neural activations and functional connectivity within the fronto-parietal regions (Cole et al., 2012; Duncan et al., 2020). In our own prior work (Etzel et al., 2020), we identified clear patterns of fronto-parietal genetic variation related to cognitive control, using a twin-based design and multivariate pattern similarity approach with the N-back task. Nevertheless, the current study departs from our prior one and from other neuroimaging work, in that those prior studies were able to directly estimate genetic heritability effects by incorporating dizygotic (fraternal) as well as monozygotic twins. In the current work, we chose to optimize our focus towards the demonstration of both domain-generality and task-specificity through the use of multiple tasks with temporally precise event-related designs. The promising results of this somewhat smaller-scale study suggest the potential of extending the sample to include both dizygotic twins and related siblings, in order to better tease apart the underlying genetic and environmental contributions that underlie individual variation in the neural mechanisms of cognitive control. Furthermore, the identification of neural pattern similarity as a key marker of genetic influences within paired individuals opens up avenues for future research utilizing a longitudinal approach, to explore the effects of developmental or interventional effects on neural similarity among these pairs. Finally, although we began an initial investigation to examine potential relationships between functional and anatomic similarity (finding none), this is an area that could be expanded upon with more sophisticated anatomical measures (e.g., voxel-based morphometry, cortical thickness, anatomic deformation, etc.). In the current study, the goal was purely to demonstrate that anatomic similarity among twins was unlikely to be the primary source of variation that accounts for the functional similarity in neural activation patterns during cognitive control.

### 4.3 Temporal specificity of twin similarity

A key feature of our approach was the use of event-related task designs to probe cognitive control demands, which enable examination of the full timecourse of pattern similarity effects throughout the trial. To our knowledge this is the first time such an approach has been utilized in conjunction with a twin-based design, particularly across multiple tasks (cf. Koten et al., 2009). The results of the timecourse analyses provided further evidence of the specificity of twin-related similarity effects within fronto-parietal regions. In particular, the peak of the twin similarity effects was observed during the same time period of task trials in which cognitive control demand peaked (as observed from univariate task-activation profiles) for 3 of the 4 tasks. Moreover, this temporal specificity was only found in the FPN parcels. It is also important to note that although there may be a relationship between the time periods of peak activation intensity in brain regions and those when twin similarity effects are most prominent, this is unlikely to be due to methodological artifact or activation differences between the two groups. In particular, the mean activation timecourses of the twin and unrelated pairs were almost identical, indicating that the twin similarity effects were not a consequence of increased twin-related activation intensity effects. These timecourse analyses also suggest a departure from the interpretations of the one prior neuroimaging study to examine twin-effects related to task events and control manipulations (Koten et al., 2009; a study using working memory load and distraction manipulations). In that study, twin effects only partially overlapped with the anatomical locations and time periods exhibiting the strongest activation effects. One potential source of this discrepancy is that the sample size and number of twin pairs in the current study is much greater (n=94, 28 twin-pairs vs. n=30, 10 twin-pairs), potentially leading to more stability in the location and estimates. It is also possible that the discrepant profile of results was due to the differences in tasks and focus of the experimental manipulations, since the prior work examined working memory load effects, whereas the current study was optimized for examination of cognitive control demands.

The utility of the time-course analyses also suggests potential directions for future work. For example, although the peak of the twin similarity effects occurred during target or probe processing, some theoretical models predict temporal shifts in peak cognitive control utilization related to shifts in cognitive control mode. In particular, the Dual Mechanisms of Control framework (Braver, 2012) suggest a distinction between proactive and reactive modes of control. In the proactive control mode, control processes are hypothesized to be sustained and anticipatory, and so likely to be engaged and even strongest prior to target / probe onsets, such as during cue or delay periods in working memory tasks, and in cued tasks such as the AX-CPT and Cued-TS. Thus, one promising research direction would be to test whether twin similarity effects shift in terms of their temporal profile under conditions that encourage proactive control utilization. A strong hypothesis would be that shifts in the time-course of similarity effects in twin pairs would be predictive of a shift in behavioral markers indicating increased utilization of proactive control.

### 4.4 Limitations

A primary limitation of the current study, alluded to previously, is that because the research design contrasted monozygotic (identical) twins and unrelated individuals, it is not possible to use these data to tease apart genetic and environmental contributions to individual differences in the neural representations (i.e., activation patterns) of cognitive control. Thus, the observed twin similarity effects cannot be entirely attributed to genetic relatedness. Because the current study aimed to serve as a first stage of investigations into the nature of individual differences in cognitive control rather than a systematic genetic modeling attempt, future large-scale neuroimaging studies comparing twins with other sets of related pairs (e.g., dizygotic twins, siblings) are needed to comprehensively evaluate the degree to which gene and environment each influences cognitive control neural activations. A second limitation of the study was that the sample was not optimized to examine gender differences in the neural representations of cognitive control, which may serve as another source of variation that impacts cognitive control function. As the focus of the current study was on twin effects, future large-scale work is needed to evaluate the extent to which gender may contribute to individual variability in cognitive control after accounting for genetic and environmental influences.

## 5. Conclusions

The current study is the first to utilize a multi-task, event-related fMRI twin-based design to systematically examine neural activation patterns related to cognitive control demands in terms of both individual variability and potentially heritable similarity effects. By leveraging our theoretically-guided task battery of cognitive control, we were able to provide promising evidence that genetic influences contribute, at least in part, to individual differences in the neural representation of cognitive control processes. The approach leveraged here, defining twin-similarity effects in terms of multivariate neural activation patterns, highlights the fine-grained nuances of cognitive control variation that are occurring at the level of neural activation patterns. Critically, this individual variation demonstrates both anatomical and temporal specificity, yet generalizability across different experimental assays of cognitive control function. In this regard, our findings offer both new insights and targets for basic and translational research aimed at understanding the neural mechanisms that give rise to individual differences in cognitive control function.

## Acknowledgments

This work was supported by NIH R37 MH066078 and R21 AT009483 to T.S.B., and F31 AT010422 to R.T.

## Author Contributions

Conceptualization, T.S.B., R.T., and J.A.E.; Investigation, R.T. and A.K.; Formal Analysis, R.T. Writing – Original Draft, R.T. and T.S.B.; Writing – Review & Editing, R.T., T.S.B., J.A.E., and A.K.; Funding Acquisition, T.S.B.

